# A conserved Lkb1 signaling axis modulates TORC2 signaling in budding yeast

**DOI:** 10.1101/293209

**Authors:** Maria Alcaide-Gavilán, Rafael Lucena, Katherine Schubert, Karen Artiles, Jessica Zapata, Douglas R. Kellogg

## Abstract

Nutrient availability, growth rate and cell size are closely linked. For example, in budding yeast, the rate of cell growth is proportional to nutrient availability, cell size is proportional to growth rate, and growth rate is proportional to cell size. Thus, cells grow slowly in poor nutrients and are nearly half the size of cells growing in rich nutrients. Moreover, large cells grow faster than small cells. A signaling network that surrounds Tor kinase complex 2 (TORC2) plays an important role in enforcing these proportional relationships. Cells that lack components of the TORC2 network fail to modulate their growth rate or size in response to changes in nutrient availability. Here, we show that budding yeast homologs of the Lkb1 tumor suppressor kinase are required for normal modulation of TORC2 signaling and in response to changes in carbon source. Lkb1 kinases activate Snf1/AMPK to initiate transcription of genes required for utilization of poor carbon sources. However, Lkb1 influences TORC2 signaling via a novel pathway that is independent of Snf1/AMPK. Of the three Lkb1 homologs in budding yeast, Elm1 plays the most important role in modulating TORC2. Elm1 activates a pair of related kinases called Gin4 and Hsl1. Previous work found that loss of Gin4 and Hsl1 causes cells to undergo unrestrained growth during a prolonged mitotic arrest, which suggests that play a role in linking cell cycle progression to cell growth. We found that Gin4 and Hsl1 also control the TORC2 network. In addition, Gin4 and Hsl1 are themselves influenced by signals from the TORC2 network, consistent with previous work showing that the TORC2 network constitutes a feedback loop. Together, the data suggest a model in which the TORC2 network sets growth rate in response to carbon source, while also relaying signals via Gin4 and Hsl1 that set the critical amount of growth required for cell cycle progression. This kind of close linkage between control of cell growth and size would suggest a simple mechanistic explanation for the proportional relationship between cell size and growth rate.

## INTRODUCTION

Growth is a defining feature of life, and always occurs at the level of cells. Mechanisms that control the rate, extent, location and timing of cell growth are responsible for the myriad forms of life that have been produced by evolution. Mechanisms that detect and limit growth during the cell cycle ultimately define the size of proliferating cells. Control of cell growth also requires mechanisms that coordinate the diverse processes of growth, including membrane growth and biogenesis, to ensure that the rate of each process is matched to the availability of building blocks.

A number of observations have pointed to close relationships between nutrient availability, growth rate, and cell size. Limiting nutrients causes reduced growth rate as well as coordinated changes in the transcription of over a thousand genes, which suggests extensive coordination of growth-related processes (Brauer *et al.* 2008). In addition, growth rate has a strong influence on cell size. Thus, cell size is proportional to growth rate, which means that slow growing cells can be nearly half the size of rapidly growing cells (Johnston *et al.* 1977; Fantes and Nurse 1977). Conversely, at least in some cases cell size influences growth rate so that large cells grow faster than small cells (Tzur *et al.* 2009; Sung *et al.* 2013; Schmoller *et al.* 2015; Leitao and Kellogg 2017). Together, these observations show that growth rate is matched to nutrient availability, cell size is matched to growth rate, and growth rate is matched to cell size. There is evidence that these relationships hold across all orders of life (Schaechter *et al.* 1958; Hirsch and Han 1969; Johnston *et al.* 1977).

In budding yeast, modulation of cell size and growth rate in response to carbon source is dependent upon a signaling network that surrounds a multi-protein kinase complex known as TOR complex 2 (TORC2) (Lucena *et al.* 2017b). TORC2 directly phosphorylates and activates a pair of partially redundant kinase paralogs called Ypk1 and Ypk2, which are the budding yeast homologs of vertebrate SGK kinases (Kamada *et al.* 2005; Niles *et al.* 2012). Full activity of Ypk1/2 also requires phosphorylation by Pkh1 and Pkh2, another pair of kinase paralogs that are the yeast homologs of vertebrate PDK1 (Casamayor *et al.* 1999). Expression of constitutively active Ypk1 rescues lethality caused by inactivation of TORC2, which suggests that Ypk1/2 are amongst the most important targets of TORC2 (Kamada *et al.* 2005; Niles *et al.* 2012). Activation of SGK kinases by TORC2 and PDK1 is conserved in vertebrates (Biondi *et al.* 2001; García-Martínez and Alessi 2008).

The TORC2-Ypk1/2 signaling axis controls production of sphingolipids and ceramides, which play roles in signaling and also serve as precursors for synthesis of structural lipids (**Figure 1A**). Ypk1/2 promote synthesis of sphingolipids by relieving inhibition of serine palmitoyltransferase, the enzyme that catalyzes the first step in sphingolipid synthesis (Breslow *et al.* 2010; Roelants *et al.* 2011). Ypk1/2 also directly phosphorylate and stimulate ceramide synthase, which builds ceramides from sphingolipid precursors (Aronova *et al.* 2008; Muir *et al.* 2014). Several observations suggest that the TORC2 network is controlled by a negative feedback loop in which the TORC2 network promotes production of ceramides, while ceramides relay signals that repress TORC2 signaling. For example, inhibition of ceramide synthesis leads to increased signaling from TORC2 to Ypk1/2 (Roelants *et al.* 2011; Berchtold *et al.* 2012; Lucena *et al.* 2017b). Conversely, addition of exogenous sphingolipids causes repression of TORC2 signaling that is dependent upon conversion of sphingolipids to ceramides (Lucena *et al.* 2017b). Feedback signaling appears to depend upon membrane trafficking events that deliver lipids to the plasma membrane, which suggests that the feedback loop monitors delivery of ceramides to the plasma membrane, rather than their synthesis at the endoplasmic reticulum (Clarke *et al.* 2017). Feedback signals are partially dependent upon Rts1, a conserved regulatory subunit of PP2A (Lucena *et al.* 2017b).

**Figure 1:**
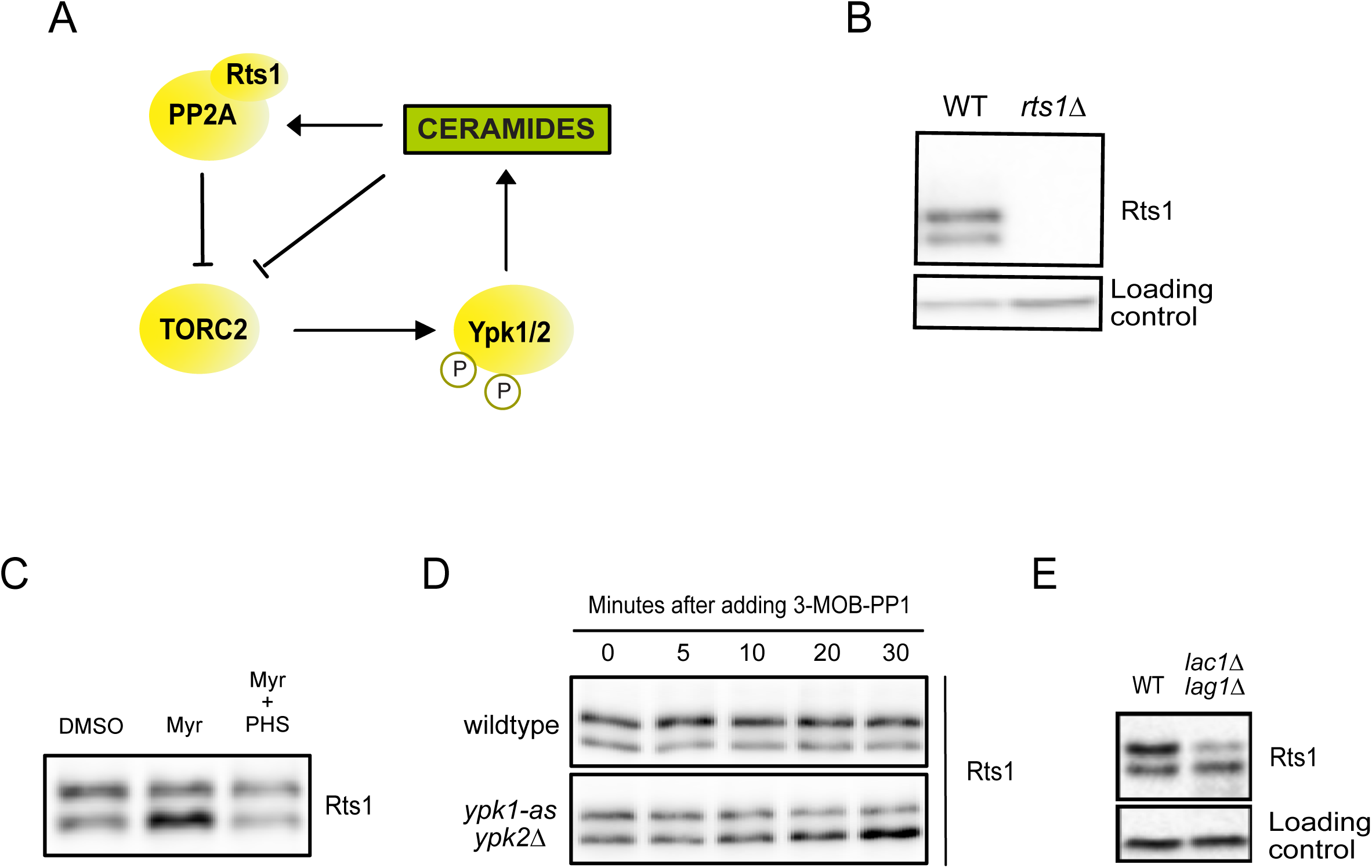
PP2A^Rts1^ is controlled by ceramide-dependent signals. (A) Wildtype and *rts1∆* cells were grown to log phase in YPD medium. Rts1 was detected by western blot using a polyclonal anti-Rts1. A background band detected by the polyclonal antibody was used as a loading control. (B) Rts1 phosphoforms were detected in log phase wildtype cells grown in the presence of 1 µM Myriocin, 20 µM of PHS or both for 90 min. Control cells were treated with dimethyl sulfoxide. (C) Wildtype and *ypk1-as ypk2∆* cells were grown to log phase in YPD medium. The adenine analog inhibitor 3-MOB-PP1 was added to both cultures at a final concentration of 50 µM to inactivate *ypk1-as*. (D) Cells of the indicated genotypes were grown to log phase in YPD and samples were taken to detect Rts1.

Diverse observations demonstrate that signals from the TORC2 feedback loop strongly influence cell growth and size (Lucena *et al.* 2017b). Shifting cells from rich to poor carbon causes a nearly two-fold decrease in cell size that is correlated with a dramatic decrease in TORC2 signaling. Decreased activity of Ypk1/2, or inactivation of ceramide synthase, cause a large reduction in cell size that is similar to the reduction caused by poor carbon. Furthermore, cells that cannot make ceramides show a complete failure in modulation of TORC2 signaling, cell size, and growth rate in response to changes in carbon source. Conversely, loss of Rts1 leads to increased TORC2 signaling due a failure in feedback signals, as well as increased cell size. Loss of Rts1 also causes a failure to properly modulate both TORC2 signaling and cell size in response to carbon source. Thus, the data suggest that the TORC2 feedback loop relays signals regarding nutrient status that help set growth rate and cell size to match nutrient availability. Genetic analysis suggests that ceramides could be the output of the network that most directly influences cell size and growth rate. However, the mechanisms and outputs of ceramide-dependent signaling are poorly understood so it remains unclear how ceramides influence cell size and growth rate.

Little is known about how nutrients influence TORC2 signaling, or how TORC2-dependent signals influence cell size. Here, we show that the yeast homolog of the Lkb1 tumor suppressor kinase is required for modulation of TORC2 signaling in response to carbon source. Nutrient modulation of TORC2 signaling further requires a set of related Lkb1-regulated kinases that were previously implicated in cell size control. Together, the data suggest a close mechanistic linkage between control of cell growth and size.

## RESULTS AND DISCUSSION

### PP2A^Rts1^ is controlled by ceramide-dependent signals

PP2A^Rts1^ relays feedback signals that influence the level of signaling in the TORC2 network (Lucena *et al.* 2017b). Previous studies suggested that ^Rts1^ is a phospho-protein (Shu *et al.* 1997), which suggested that feedback signals could be relayed via phosphorylation of Rts1. To investigate further, we raised an antibody that recognizes Rts1. The antibody recognized two forms of Rts1 in rapidly growing cells (**Figure 1B**). Treatment with phosphatase caused the forms to collapse into one band, confirming that ^Rts1^ is phosphorylated (**Supplementary Figure 1A**).

To test whether Rts1 phosphorylation could play a role in relaying feedback signals, we first determined whether inhibition of ceramide synthesis causes changes in Rts1 phosphorylation. Ceramides are made from sphingolipid precursors that are synthesized by the enzyme serine palmitoyltransferase (SPT), which can be inhibited with myriocin (Miyake *et al.* 1995; Dickson 2008; Breslow and Weissman 2010). Addition of myriocin to cells growing in rich carbon caused a partial loss of Rts1 phosphorylation. The loss was rescued by addition of the sphingolipid phytosphingosine (PHS) to the cell culture (**Figure 1C**).

The kinases Ypk1/2 promote synthesis of ceramides in the feedback loop by directly phosphorylating ceramide synthase (Muir *et al.* 2014). To test whether Ypk1/2 are required for normal control of PP2A^Rts1^ phosphorylation, we utilized an analog-sensitive allele of Ypk1 in a *ypk2∆* background (*ypk1-as ypk2∆*) (Sun *et al.* 2012). Inhibition of Ypk1/2 in cells growing in rich carbon caused a loss of ^Rts1^ phosphorylation that was similar to the loss caused by myriocin (**Figure 1D**).

We next used mutants and inhibitors that block synthesis of specific sphingolipids to determine which products of sphingolipid processing are required for normal phosphorylation of Rts1. Ceramide synthase is encoded by redundant paralogs called *LAC1* and *LAG1* (Guillas *et al.* 2001; Schorling *et al.* 2001). ^Rts1^ phosphorylation was largely lost in *lac1∆ lag1∆* cells (**Figure 1E**). Mutants or inhibitors that block all other known sphingolipid processing steps had no effect on ^Rts1^ phosphorylation (**Supplemental Figure 1B**).

Together, these observations demonstrate that ceramide-dependent signals control ^Rts1^ phosphorylation, consistent with previous genetic data that suggested that PP2A^Rts1^ relays ceramide-dependent feedback signals in the TORC2 network (Lucena *et al.* 2017b). Neither Ypk1 or Pkh1 could phosphorylate ^Rts1^ in vitro, which suggests that they act indirectly. Moreover, kinases in the TORC2 network are unlikely to directly phosphorylate ^Rts1^ because loss of ceramides is thought to cause increased activity of TORC2 network kinases, yet causes decreased ^Rts1^ phosphorylation (Roelants *et al.* 2011; Berchtold *et al.* 2012; Lucena *et al.* 2017b). Despite extensive efforts, the kinase or kinases responsible for direct phosphorylation of Rts1 in response to ceramide-dependent signals remain unknown. A screen of all viable kinases deletions did not identify additional kinases required for phosphorylation of Rts1. Similarly, a screen of conditional alleles covering essential kinases, including redundant paralogs, identified only Ypk1/2 and Pkh1/2. A potential explanation is that ^Rts1^ phosphorylation in this context is carried out by multiple redundant kinases.

### Rts1 phosphorylation and TORC2 signaling are correlated with growth rate during the cell cycle

We previously proposed that the level of signaling in the TORC2 feedback loop plays a role in setting growth rate (Lucena *et al.* 2017b). This model predicts that TORC2 signaling should be proportional to growth rate. Moreover, if phosphorylation of ^Rts1^ helps relay feedback signals that set the level of TORC2 signaling, then ^Rts1^ phosphorylation should also be proportional to growth rate. To test these predictions, we took advantage of the fact that the rate of growth varies during the cell cycle. Cells grow slowly during interphase, but the rate of growth increases approximately 3-fold when cells enter mitosis (Goranov *et al.* 2009; Leitao and Kellogg 2017). The rapid mitotic growth phase accounts for most of the volume of a budding yeast cell (Leitao and Kellogg 2017). To test whether TORC2 signaling and ^Rts1^ phosphorylation are correlated with growth rate, we released small G1 cells isolated from centrifugal elutriation and assayed TORC2 activity and ^Rts1^ phosphorylation. TORC2 activity was assayed with a phosphospecific antibody that recognizes a TORC2 site present on Ypk1 and Ypk2 (Niles *et al.* 2012). The same samples were probed for the mitotic cyclin Clb2 to provide a marker for mitosis (**Figure 2A**). TORC2 activity increased approximately 3-fold in mitosis (**Figure 2B**). In addition, ^Rts1^ was predominantly in the lower form early in G1 phase, but shifted more to the slower mobility form during mitosis (**Figure 2C**). Thus, growth rate during the cell cycle is proportional to both ^Rts1^ phosphorylation and TORC2 signaling, further demonstrating a link between TORC2 signaling and growth rate.

**Figure 2:**
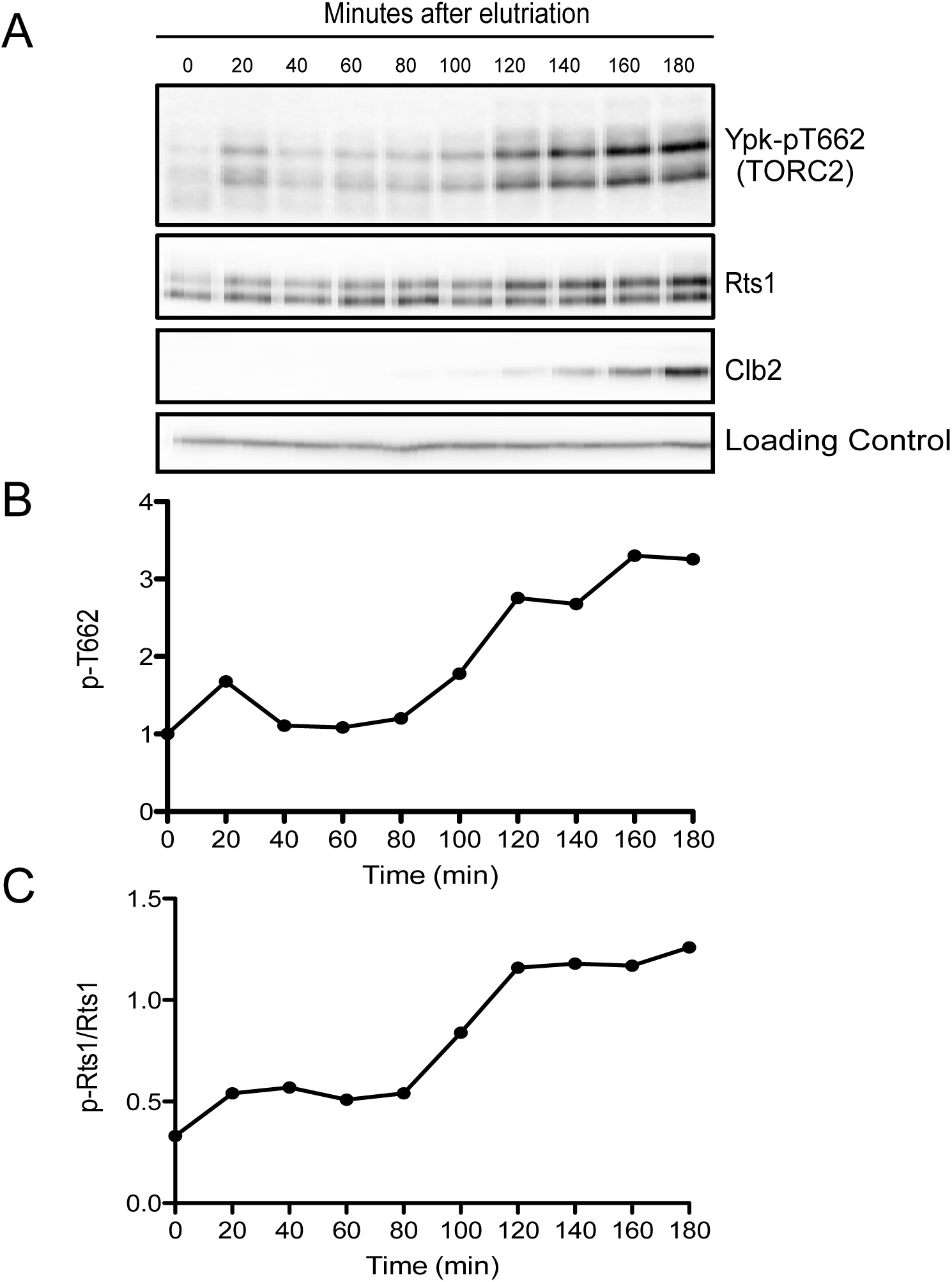
Rts1 phosphorylation and TORC2 signaling are correlated with growth rate during the cell cycle. (A) Wildtype cells were grown to log phase in YPG/E medium. Small unbudded cells in early G1 phase were isolated by centrifugal elutriation and released into YPD medium at 25^º^C. Samples were taken every 20 minutes and Ypk-pT662, Rts1 and Clb2 were analyzed by western blot. (B) Quantification of p-T662 in the samples shown in panel A.(C) Quantification of the phosphorylated form of Rts1 in the samples shown in panel A. The ratio of the phosphorylated form of Rts1 over the unphosphorylated form is shown for each time point.

### PP2A^Rts1^ is controlled by nutrients

The level of signaling in the TORC2 network is proportional to the growth rate set by nutrient availability (Lucena *et al.* 2017b). Thus, when cells are shifted from rich to poor carbon, there is an immediate and dramatic reduction in TORC2 signaling. Over several hours, TORC2 signaling recovers as cells adapt to poor carbon, but remains depressed relative to cells in rich carbon. PP2A^Rts1^ is not required for the immediate response to poor carbon. However, in cells that have undergone adaptation to poor carbon, TORC2 signaling is elevated in *rts1∆* cells relative to wild type cells. Therefore, we next tested whether Rts1 phosphorylation is influenced by nutrient-dependent signals. We first assayed Rts1 phosphorylation in cells growing in carbon sources of varying quality. Yeast cells utilize glucose most efficiently. Galactose is an intermediate quality carbon source, while glycerol and ethanol are poor carbon sources. A fraction of Rts1 was hyperphosphorylated in cells growing in poor carbon sources (**Figure 3A**). We refer to the phosphorylated form of Rts1 observed in rich carbon as an intermediate phosphorylated form (marked with a single asterisk in **Figure 3A**) and the slower mobility forms observed in poor carbon as hyperphosphorylated Rts1 (marked with two asterisks in **Figure 3A**).

**Figure 3:**
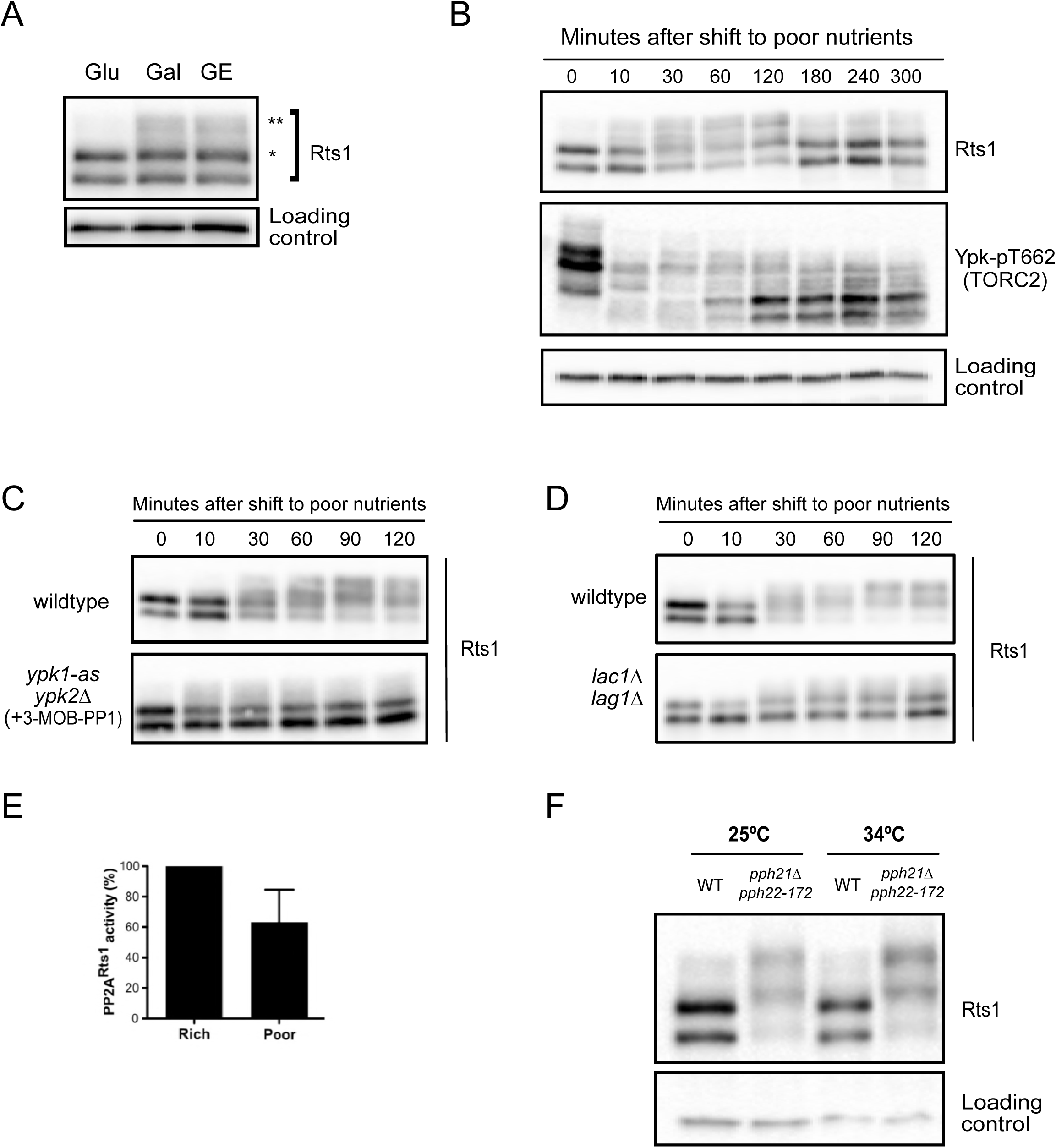
PP2A^Rts1^ is controlled by nutrients. (A) Wildtype cells were grown to early log phase in YEP media containing glucose (Glu), galactose (Gal) or glycerol/ethanol (GE) and Rts1 was assayed by western blot. A single asterisk marks a phosphorylated form of Rts1; two asterisks are used to mark a hyperphosphorylated form of Rts1. (B) Wildtype cells were grown in YPD medium to log phase and were then shifted to YPG/E medium. Cells were collected up to 300 minutes and Rts1 and Ypk-pT662 were analyzed by western blot. (C) Cells of the indicated genotypes were treated shifted from YPD to YPG/E medium. 50 µM 3-MOB-PP1 was added to both cultures immediately after the cells were shifted to YPG/E. Rts1 was detected by western blot. (D) Cells of the indicated genotypes were treated shifted from YPD to YPG/E medium. Rts1 was detected by western blot. (E) Rts1-associated phosphatase activity was assayed in log phase wild type cells and in wild type cells 90 minutes after a shift to YPG/E media. The activity in YPG/E medium (poor) was plotted as a percentage of the activity in YPD medium (Rich). (F) Wildtype and *pph21∆ pph22-172* cells were grown to early log phase at 25^º^C and then shifted to 34^º^C for 15 minutes. Rts1 was detected by western blot.

We next analyzed the effects of an acute shift from rich to poor carbon. Within minutes of a shift to poor carbon, Rts1 underwent extensive hyperphosphorylation that persisted for several hours (**Figure 3B**). Hyperphosphorylation of Rts1 was reduced as cells adapted to the new carbon source, but remained elevated relative to cells growing in rich carbon.

Two observations suggest that the initial rapid hyperphosphorylation of Rts1 in poor carbon is not a cause of the acute reduction in TORC2 signaling. First, the abrupt decrease in TORC signaling caused by a shift to poor carbon occurred more rapidly than hyperphosphorylation of Rts1. Second, Rts1 is not required for the abrupt reduction in TORC2 activity (Lucena *et al.* 2017b). Rather, hyperphosphorylation of Rts1 in response to poor carbon may be due to reduced feedback signaling in the TORC2 network. Consistent with this, we found that inhibition of Ypk1/2 or inactivation of ceramide synthase largely eliminated hyperphosphorylation of Rts1 in response to poor carbon (**Figures 3C, D**). The response of Rts1 to poor carbon was identical in the A364A strain background, but was substantially different in the S288C strain background (**Supplementary Figure S2**).

We next tested whether PP2A activity is correlated with Rts1 phosphorylation. To do this, we immunoprecipitated Rts1 from cells growing in rich or poor carbon and measured associated phosphatase activity. PP2A activity associated with Rts1 was reduced in cells growing in poor carbon, when Rts1 was hyperphosphorylated (**Figure 3E**). A previous study found that Rts1 dissociates from PP2A when cells are grown in poor carbon, which could explain the reduction in Rts1-associated phosphatase activity in those conditions (Castermans *et al.* 2012). Further evidence for dissociation of the PP2A^Rts1^ complex in poor carbon came from analysis of Rts1 phosphorylation in response to inactivation of PP2A catalytic subunits. Since Rts1 is bound to PP2A, we hypothesized that it could be subject to PP2A-dependent dephosphorylation. To test this, we assayed Rts1 phosphorylation in cells dependent upon a temperature-sensitive allele of one of the redundant PP2A catalytic subunits (*pph21∆ pph22-172*). Wildtype and *pph21∆ pph22-172* cells were shifted to the restrictive temperature and Rts1 phosphorylation was analyzed by western blot. Highly hyperphosphorylated forms of Rts1 were observed even at the permissive temperature, consistent with the idea that Rts1 phosphorylation is opposed by PP2A (**Figure 3F**). Thus, decreased PP2A activity causes hyperphosphorylation of Rts1 that is similar to hyperphosphorylation of Rts1 caused by a shift to poor carbon, consistent with a model in which poor nutrients cause dissociation of the PP2A^Rts1^ complex, leading to hyperphosphorylation of Rts1.

Together, the data suggest that a shift from rich to poor carbon triggers changes in ceramide-dependent signaling that lead to dissociation of the PP2A^Rts1^ complex and reduced PP2A^Rts1^ activity. These observations point to a paradox: a complete loss of PP2A^Rts1^, as seen in *rts1∆* cells, causes increased cell size in both rich and poor carbon. In contrast, poor nutrients appear to cause a partial loss of PP2A^Rts1^ activity, yet cause decreased cell size. A potential explanation is that Rts1 dissociates from PP2A in poor carbon and then carries out PP2A-independent functions. A previous study suggested that Rts1 associates with Glc7, the budding yeast homolog of the PP1 phosphatase, and that association of Rts1 with Glc7 increases in poor carbon (Castermans *et al.* 2012). Thus, the PP2A^Rts1^ complex could dominate in rich carbon, while a Glc7^Rts1^ complex could play a role in the response to poor carbon, including a role in reducing cell size.

### Budding yeast homologs of vertebrate Lkb1 kinase modulate TORC2 signaling

We next searched for proteins that relay signals regarding carbon source to the TORC2 network. The kinases Elm1, Tos3 and Sak1 were strong candidates. Together, these kinases are related to vertebrate Lkb1, an important tumor suppressor, and their functions can be replaced by Lkb1, indicating that they are true functional homologs of Lkb1 (Hong *et al.* 2003; Sutherland *et al.* 2003; Woods *et al.* 2003). An important function of Lkb1-related kinases is to activate AMPK, which initiates transcription of hundreds of genes required for utilization of poor energy sources. The budding yeast AMPK homolog is referred to as Snf1. Loss of Snf1, or all three of the Snf1-activating kinases, causes a failure to proliferate on poor carbon sources. Elm1 is of particular interest because it was previously implicated in control of cell growth and size (Blacketer *et al.* 1993; Sreenivasan and Kellogg 1999). For example, loss of Elm1 causes a prolonged delay in mitosis. During the delay, polar growth of the daughter bud continues, leading to growth of abnormally large cells. Loss of vertebrate Lkb1 also causes an increase in cell size (Faubert *et al.* 2014). Furthermore, *elm1∆* shows genetic interactions with G1 cyclins that suggest that it could function with Cln3, a G1 cyclin required for control of cell size in G1 phase (Sreenivasan *et al.* 2003). Although Tos3 and Sak1 share functions with Elm1 in phosphorylation of Snf1, they do not appear to be required for control of cell size, and they are not known to show genetic interactions with late G1 cyclins.

We first tested whether Elm1 is required for normal control of the TORC2 network. In rich carbon, *elm1∆* caused a decrease in TORC2-dependent phosphorylation of Ypk1/2, but had no effect on Rts1 phosphorylation (**Figure 4A**). In a shift from rich to poor carbon, *elm1∆* caused reduced hyperphosphorylation of Rts1, as well as a failure in modulation of TORC2-dependent phosphorylation of Ypk1/2 (**Figure 4B**). Combined loss of Elm1, Tos3 and Sak1 caused a complete failure in hyperphosphorylation of Rts1 in response to poor carbon (**Figure 4B**). In contrast, *snf1∆* had no effect on modulation of TORC2 signaling in response to carbon source, which indicates that Elm1/Tos3/Sak1 influence TORC2 signaling via a novel pathway (**Figure 4C**). Growth in poor carbon did not reduce the abnormally large size of *elm1∆* cells, which suggests a failure in nutrient modulation of cell size (**Figure S3**).

**Figure 4:**
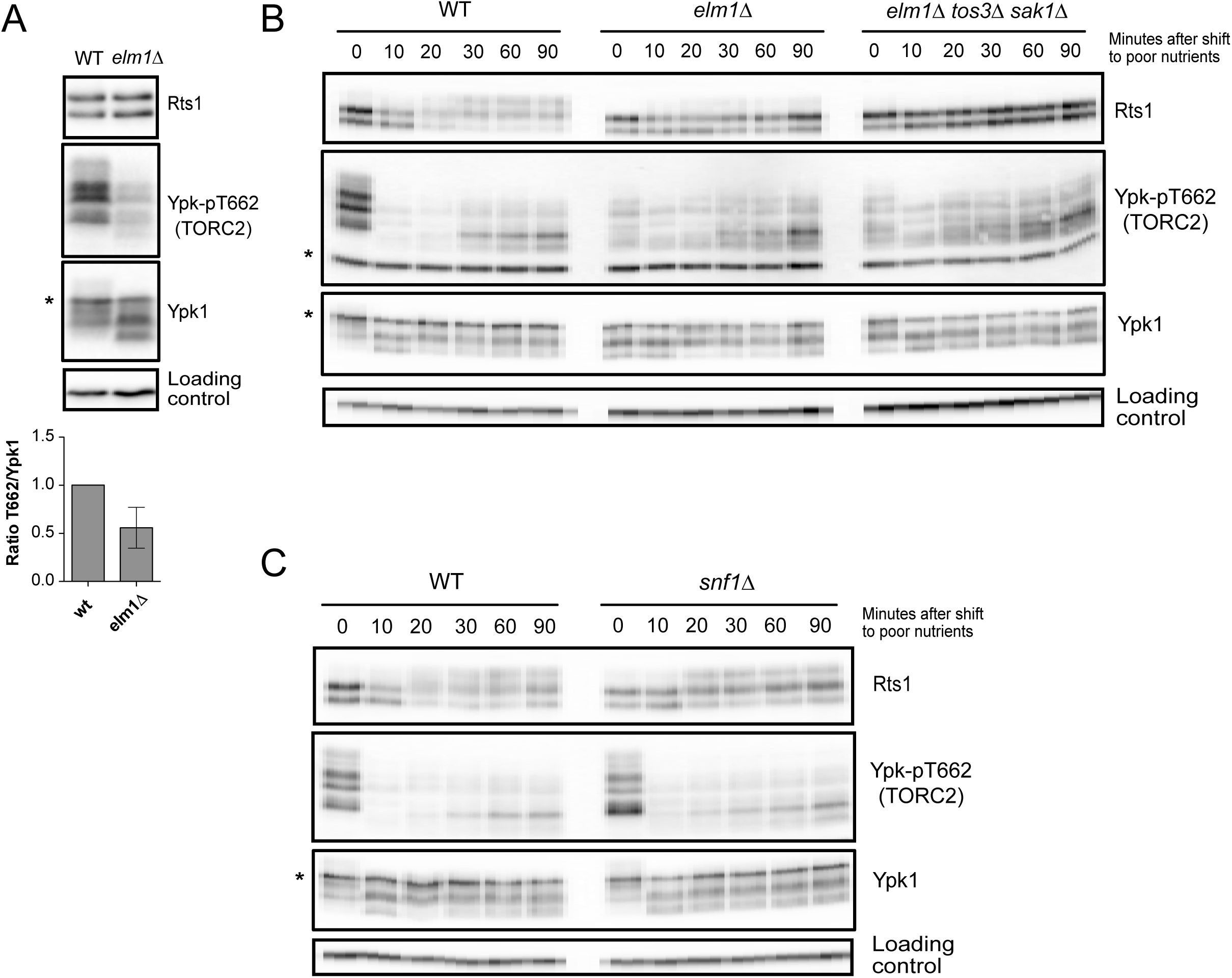
Budding yeast homologs of vertebrate Lkb1 kinase modulate TORC2 signaling. Cells of the indicated genotypes were grown to log phase in YPD medium. Rts1, Ypk-pT662 and Ypk1 were analyzed by western blot. Quantification of the Ypk-pT662 signal over total Ypk1 protein is shown. Error bars represent the standard deviation of the mean of 3 biological replicates. (B, C) Cells of the indicated genotype were grown to log phase in YPD medium and were then shifted to YPG/E medium. Rts1, Ypk-pT662 and Ypk1 were detected by western blot. An asterisk indicates a background band.

### Gin4-related kinases are required for nutrient modulation of TORC2 signaling

Elm1 controls the activities of three related kinases called Gin4, Kcc4 and Hsl1, which we refer to as Gin4-related kinases. It is thought that Elm1 directly phosphorylates the Gin4-related kinases to stimulate their activity, and combined loss of Gin4 and Hsl1 causes a phenotype similar to *elm1∆* (Barral *et al.* 1999; Asano *et al.* 2006; Szkotnicki *et al.* 2008). Kcc4 does not appear to play a major role. Fission yeast homologs of Gin4 and Hsl1 are required for normal control of cell size in rich nutrients and for nutrient modulation of cell size (Young and Fantes 1987). An important and conserved function of the Gin4-related kinases is to promote progression through mitosis via inhibition of the Wee1 kinase, an inhibitor of mitotic progression. Loss of Wee1 causes premature progression through mitosis and reduced cell size in both budding yeast and fission yeast, and the large size of yeast cells that lack Gin4-related kinases is partially rescued by inactivation of Wee1 (Ma *et al.* 1996; Barral *et al.* 1999; Jorgensen *et al.* 2002; Harvey and Kellogg 2003; Harvey *et al.* 2005). However, the mechanisms by which Gin4-related kinases inhibit Wee1 are poorly understood.

To investigate further, we tested whether loss of the Gin4-related kinases caused effects on TORC2 signaling. In cells growing in rich carbon, loss of individual Gin4-related kinases caused little or no effect on TORC2 signaling, but combined loss of all three caused a reduction in TORC2 signaling similar to the reduction caused by *elm1∆*. (**Figures 5A).** In a shift from rich to poor carbon, cells lacking all three Gin4-related kinases showed reduced hyperphosphorylation of Rts1, which can be seen as an increase in the fraction of Rts1 found in the dephosphorylated and intermediate phosphorylated forms. Loss of the Gin4-related kinases also caused defective modulation of TORC2 signaling in response to poor carbon (**Figures 5B**).

**Figure 5:**
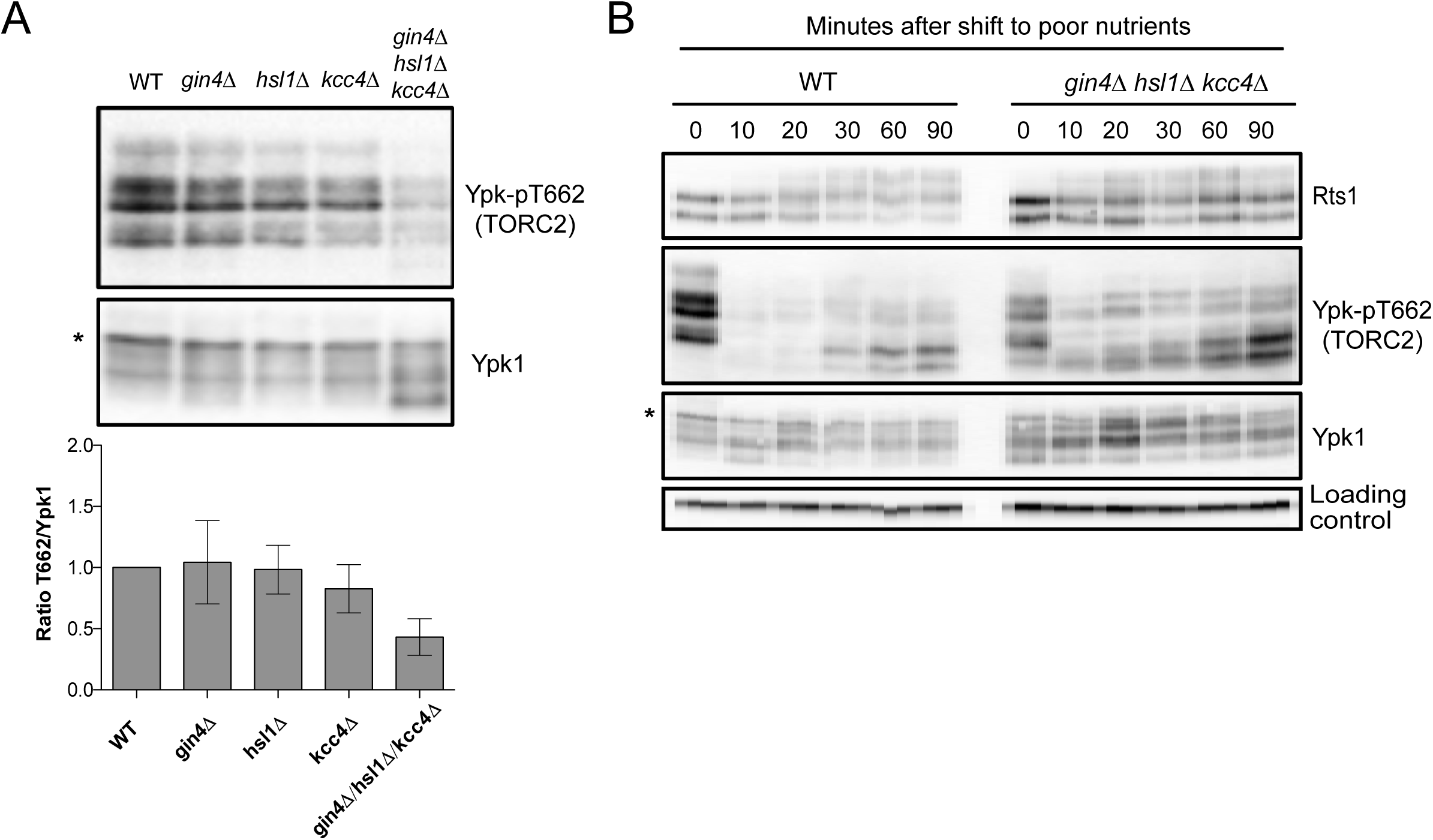
Gin4-related kinases are required for nutrient modulation of TORC2 signaling. (A) Cells of the indicated genotypes were grown to log phase in YPD medium. Rts1, Ypk-pT662 and Ypk1 were detected by western blot. The Ypk-pT662 signal was quantified as a ratio of the pT662 signal over the Ypk1 signal. Error bars represent the standard deviation of the mean of 3 biological replicates. (B) Wildtype and Gin4-related kinase mutants were grown in YPD medium and were then shifted to YPG/E medium. Rts1, Ypk-pT662 and Ypk1 were detected by western blot. An asterisk indicates a background band.

We also noticed that loss of Elm1 or the Gin4-related kinases caused increased electrophoretic mobility of Ypk1 (bottom panels in **Figures 4A and 5A**). Previous work showed that shifts in the electrophoretic mobility of Ypk1 are due to phosphorylation of Ypk1 by redundant kinase paralogs Fpk1 and Fpk2, which play poorly understood roles in controlling Ypk1/2 (Roelants *et al.* 2010). The data suggest that Fpk1/2 have reduced activity in cells that lack Elm1 or the Gin4-related kinases.

The effect of *elm1∆* on Rts1 hyperphosphorylation in poor carbon appeared to be stronger than the effect caused by loss of all three Gin4-related kinases (compare **Figures 4B and 5B**). This observation suggests that Elm1 may influence Rts1 phosphorylation via multiple pathways, including one that that is independent of the Gin4-related kinases.

### Elm1 and the Gin4-related kinases do not influence TORC2 signaling solely via Rts1

We next used epistasis analysis to investigate how Elm1 and the Gin4-related kinases influence TORC2 signaling. We discovered that increased TORC2 signaling in *rts1∆* cells is dependent upon Elm1 and the Gin4-related kinases (**Figures 6A, B**). This observation rules out a model in which Elm1 and the Gin4-related kinases influence TORC2 signaling upstream of Rts1. Rather, the data suggest that Rts1 inhibits Elm1-dependent signaling to the TORC2 network. In previous work, we found that *rts1∆* causes hyperphosphorylation of Mss4, an important upstream activator of the TORC2 network (Lucena *et al.* 2017b). Hyperphosphorylation of Mss4 is thought to lead to its increased recruitment to the plasma membrane, where it stimulates TORC2 signaling. Here, we found that *elm1∆* does not cause effects on Mss4 phosphorylation that can be detected via electrophoretic mobility shifts (**Figure 6C**). This observation suggests that Elm1 and the Gin4-related kinases could influence TORC2 signaling via a mechanism that is independent of Rts1-dependent control of Mss4 phosphorylation.

**Figure 6:**
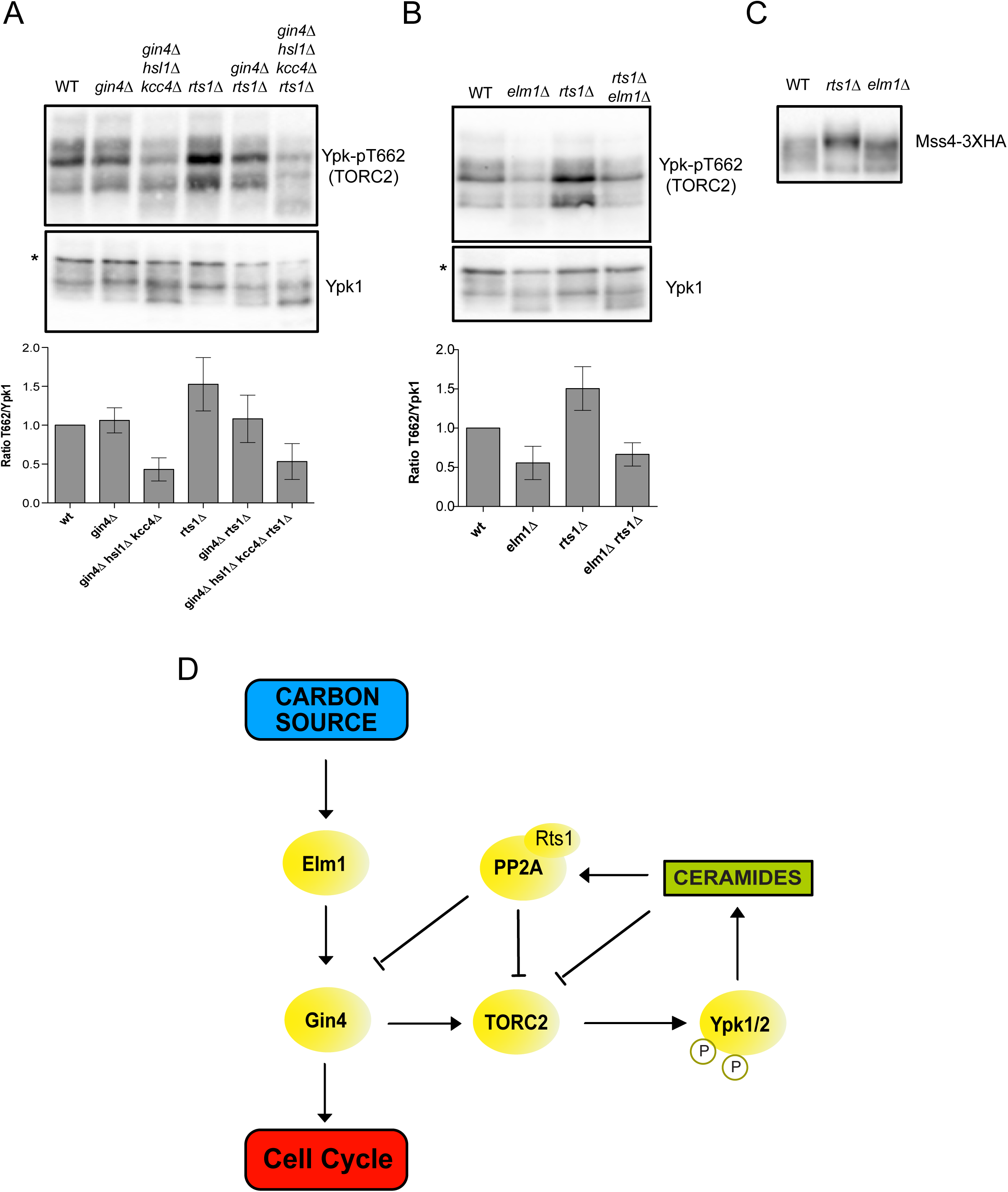
Elm1 and the Gin4-related kinases do not influence TORC2 signaling solely via Rts1. (A) Cells of the indicated genotypes were grown to log phase in YPD medium. Mss4-3XHA was detected by western blot. (B, C) Cells of the indicated genotypes were grown to log phase in YPD medium. The Ypk-pT662 signal was quantified as a ratio of the pT662 signal over the Ypk1 signal. Error bars represent the standard deviation of the mean of 3 biological replicates. (D) A proposed model that could explain the signals that influence cell growth and size via the TORC2 network. An asterisk indicates a background band.

Together, the data provide new insights into signals that influence cell growth and size via the TORC2 network. A model that could explain the data is shown in **Figure 6D**. We propose that the Lkb1-related kinases in yeast relay signals regarding carbon source to the TORC2 network. Of the Lkb1-related kinases, Elm1 plays a dominant role, as loss of Elm1 alone caused decreased TORC2 signaling as well as a failure in modulation of TORC2 signaling in response to carbon source. Previous work suggested that Elm1 activates the Gin4-related kinases in a pathway that controls Wee1 (Asano *et al.* 2006; Szkotnicki *et al.* 2008). Here, we found that loss of the Gin4-related kinases causes effects on TORC2 signaling that are similar to those caused by loss of Elm1, which suggests that Elm1 also controls the Gin4-related kinases in the context of TORC2 signaling. Cells that lack Elm1 or the Gin4-related kinases show no evidence of nutrient modulation of cell size, which provides further evidence that they influence TORC2 signaling. The Lkb1-related kinases influence TORC2 signaling independently of Snf1/AMPK, which points to the existence of a surprising new pathway that links Lkb1-related kinases to TORC2 signaling.

We further discovered that signals from the TORC2 network control phosphorylation of Rts1. In cells growing in rich nutrients, Rts1 is pushed towards an intermediate phosphorylation state during mitosis that is positively correlated with high levels of TORC2 signaling and high growth rate. In cells growing in poor carbon, Rts1 undergoes hyperphosphorylation that is correlated with slow growth rate and reduced TORC2 signaling. In both contexts, normal phosphorylation of Rts1 is dependent upon ceramides. Since ceramide production is controlled by TORC2-dependent signals, this observation strengthens previous genetic data that suggested a role for Rts1 in a ceramide-dependent feedback loop that negatively regulates TORC2 signaling (Lucena *et al.* 2017b). Rts1 hyperphosphorylation in poor carbon is correlated with a decrease in Rts1-associated phosphatase activity, consistent with a previous study that suggested that a fraction of Rts1 dissociates from PP2A in poor carbon (Castermans *et al.* 2012). A previous study suggested that the Rts1 that dissociates from PP2A in poor carbon could have PP2A-independent functions. It is not yet known whether hyperphosphorylation of Rts1 is a cause or consequence of dissociation from the PP2A complex.

Several observations suggest that the Gin4-related kinases are influenced by Rts1-dependent feedback signals from the TORC2 network. For example, proteome-wide mass spectrometry studies have found that all three of the Gin4-related kinases are strongly hyperphosphorylated on multiple sites in *rts1∆* cells, and that synthesis of ceramides influences Gin4 phosphorylation (Zapata *et al.* 2014; Lebesgue *et al.* 2017). In addition, the increased TORC2 signaling caused by *rts1∆* is dependent upon the Gin4-related kinases. Together, these observations suggest that the Gin4-related kinases influence TORC2 signaling, while TORC2 signaling influences the Gin4-related kinases. This connection is particularly intriguing because previous studies have shown that the Gin4-related kinases are required for control of cell size in both budding yeast and fission yeast (Young and Fantes 1987; Ma *et al.* 1996; Altman and Kellogg 1997). The Gin4-related kinases control the Wee1 kinase, which is thought to influence cell size by ensuring that progression through mitosis occurs only when sufficient growth has occurred (Nurse 1975; Ma *et al.* 1996; Shulewitz *et al.* 1999; Sreenivasan and Kellogg 1999; Longtine *et al.* 2000). Thus, the discovery that the Gin4-related kinases are influenced by TORC2 signaling could provide an explanation for the previous observation that TORC2 signaling influences cell size (Lucena *et al.* 2017b). Furthermore, the discovery that the Gin4-related kinases are embedded in the TORC2 feedback loop and also control Wee1 establishes further connections between cell growth control and cell size control. We propose that the Gin4-related kinases provide a readout of the level of signaling in the TORC2 feedback loop that is used to set the threshold amount of growth required for progression through mitosis. This kind of model would provide a simple mechanistic explanation for the close relationship between cell size and growth rate. Moreover, since cell size can be viewed as an outcome of growth control, it would make sense that growth control and size control evolved together and are mechanistically linked. Testing this model will require a better understanding of the mechanisms by which Elm1 and the Gin4-related kinases influence both TORC2 signaling and cell cycle progression.

## MATERIALS AND METHODS

### Yeast strains and media

All strains are in the W303 background (*leu2-3,112 ura3-1 can1-100 ade2-1 his3-11,15 trp1-1 GAL+ ssd1-d2*), with the exception of strain DDY903 used in Figure S2, which is in the S288C background (*his3-∆200, ura3-52, lys2-801, leu2-3, 112,*). Table S1 shows additional genetic features. One-step PCR-based gene replacement was used for making deletions and adding epitope tags at the endogenous locus (Longtine, 1998; Janke et al., 2004). Cells were grown in YP medium (1% yeast extract, 2% peptone, 40 mg/L adenine) supplemented with 2% dextrose (YPD), 2% galactose (YPGal), or 2% glycerol/ethanol (YPG/E). For experiments using analog-sensitive alleles, cells were grown in YPD medium without supplemental adenine. For nutrient shifts, cells were grown in YPD medium overnight to log phase. Cells were then washed 3 times with YPG/E medium and resuspended in YPG/E medium.

Myriocin (Sigma) was dissolved in 100% methanol to make a 500 ng/ml stock solution. Phytosphingosine (Avanti Polar lipids) was dissolved in 100% ethanol to make a 10 mM stock solution. Aureobasidin A (Tanaka Clontech) was dissolved in methanol to make a 5 mg/ml stock solution. The same volume of methanol or ethanol was added in control experiments.

### Production of polyclonal Rts1 antibody

An antibody that recognizes Rts1 was generated by immunizing rabbits with a fusion protein expressed in bacteria. A plasmid expressing full length Rts1 was constructed with the Gateway cloning system. Briefly, a PCR product that includes the full-length open reading frame for Rts1 was cloned into the entry vector pDONR221. The resulting donor plasmid was used to generate a plasmid that expresses Rts1 fused at its N-terminus to 6XHis-TEV, using expression vector pDEST17 (6XHis). The 6XHis-TEV-Rts1 fusion was expressed in BL21 cells and purified in the presence of 2M urea using standard procedures, yielding 10 mg from 4 L of bacterial culture. A milligram of the purified protein was used to immunize a rabbit. The 6XHis-Rts1 fusion protein was coupled to Affigel 10 (Bio-Rad) to create an affinity column for purification of the antibody.

### Western blotting

For western blots using cells growing in early log phase, cultures were grown overnight at 25^º^C to an OD_600_ of less than 0.8. After adjusting optical densities to normalize protein loading, 1.6-ml samples were collected and centrifuged at 13,000 rpm for 30 s. The supernatant was removed and 250 µl of glass beads were added before freezing in liquid nitrogen.

To analyze cells shifted from rich to poor nutrients, cultures were grown in YPD overnight at 25^º^C to an OD_600_ of less than 0.8. After adjusting optical densities to normalize protein loading, cells were washed three times with a large volume of YPG/E medium and then incubated at 30^º^C in YPG/E for the time course. 1.6-ml samples were collected at each time point.

Cells were lysed into 140 μl of sample buffer (65 mM Tris-HCl, pH 6.8, 3% SDS, 10% glycerol, 50 mM NaF, 100 mM β -glycerophosphate, 5% 2-mercaptoethanol, and bromophenol blue). PMSF was added to the sample buffer to 2 mM immediately before use. Cells were lysed in a Mini-bead-beater 16 (BioSpec) at top speed for 2 min. The samples were removed and centrifuged for 15 s at 14,000 rpm in a microfuge and placed in boiling water for 5 min. After boiling, the samples were centrifuged for 5 min at 14,000 rpm and loaded on an SDS polyacrylamide gel.

Samples were analyzed by western blot as previously described (Harvey *et al.* 2011). SDS-PAGE gels were run at a constant current of 20 mA and electrophoresis was performed on gels containing 10% polyacrylamide and 0.13% bis-acrylamide. Proteins were transferred to nitrocellulose using a Trans-Blot Turbo system (Bio-Rad Laboratories). Blots were probed with primary antibody overnight at 4°C. Proteins tagged with the HA epitope were detected with the 12CA5 anti-HA monoclonal antibody (gift of John Tamkun). Rabbit anti-phospho-T662 antibody (gift of Ted Powers, University of California, Davis) was used to detect TORC2-dependent phosphorylation of YPK1/2 at a dilution of 1:20,000 in TBST (10 mM Tris-Cl, pH 7.5, 100 mM NaCl, and 0.1% Tween 20) containing 3% Milk. Total Ypk1 was detected using anti-Ypk1 antibody (Santa Cruz Biotechnology, catalog number sc-12051) at a dilution of 1:2000.

All blots were probed with an HRP-conjugated donkey anti-rabbit secondary antibody (GE Healthcare, catalog number NA934V) or HRP-conjugated donkey anti– mouse antibody (GE Healthcare, catalog number NXA931) or HRP-conjugated donkey anti-goat (Santa Cruz Biotechnology, catalog number sc-2020) for 45–90 min at room temperature. Secondary antibodies were detected via chemiluminescence with Advansta ECL reagents.

Densitometric quantification of western blot signals was performed using ImageJ (Scheider CA et al., 2012). Quantification of Ypk-pT662 phosphorylation was calculated as the ratio of the phospho-specific signal over the total Ypk1 protein signal, with wild-type signal normalized to a value 1. In Figure 2A, the signal was normalized over the loading control band. Phospho-Rts1 was quantified using Image Lab (BioRad). At least three biological replicates were analyzed and averaged to obtain quantitative information.

### Synchronization by centrifugal elutriation

Cells were grown overnight at 25^°^C in YPG/E medium to increase the fraction of very small unbudded cells. Centrifugal elutriation was performed as previously described (Futcher 1999; McCusker *et al.* 2012). Briefly, cells were elutriated at 4^°^C at a speed of 2,800 rpm in a Beckman Coulter J6-MI centrifuge with a JE-5.0 rotor. Small unbudded cells were released into fresh YPD or YPG/E media at 25^°^C and samples were taken at 10-min intervals.

### Lambda phosphatase treatment

Lambda phosphatase treatment of Rts1 in cell extracts was carried out as previously described (Lucena *et al.* 2017a).

### Rts1-associated phosphatase activity assay

To create a substrate for phosphatase assays, 5 µg of MBP (Sigma) were incubated with 6 µl of 10X PK buffer (50 mM Tris-HCl, 10 mM MgCl_2_, 0.1 mM EDTA, 2mM DTT, 0.01% Brij 35, pH 7.5), 200 µM ATP, 2 µl ^32^P-ATP and 2 µl PKA (New England BioLabs) in a total volume of 60 µl. The reaction was incubated for 5 h at 30ºC, TCA precipitated and resuspended in phosphatase buffer (50 mM Tris-HCl pH 7.5, 5% glycerol, 0.2% Tween 20, 0.5 mM DTT).

Rts1-associated phosphatase activity was assayed by release of ^32^P-inorganic phosphate from ^32^P-MBP. 50 ml of wildtype cells were grown in YPD, washed and split to YPG/E or YPD for 90 minutes. Cell extracts made from wild type or *rts1∆* control cells were made by adding 300µl of acid-washed glass beads to frozen cell pellets followed by 500 µl of lysis buffer (50 mM HEPES-KOH, pH 7.6, 500 mM KCl, 100 mM β-glycerol phosphate, 1 mM MgCl_2_, 1 mM EGTA, 5% glycerol, 0.25% Tween 20, and 1 mM PMSF). The tubes were immediately placed into a Mini-bead-beater 16 (Biospec), shaken at top speed for 1 min and placed in an ice-water bath for 1 min. The tubes were centrifuged 1 min at 14,000 rpm and 250 µl of the supernatant was transferred to a new 1.5-ml tube and replaced with 250 µl of lysis buffer. The tubes were beaten again for 1 min, 250 µl of the supernatant was removed, pooled with the first supernatant, and centrifuged at 14,000 rpm in a microcentrifuge for 10 min at 4 ^°^C. The supernatant was added to 100 µl of protein A beads loaded with 15 µg of anti-Rts1 and equilibrated in lysis buffer. The tubes were rotated gently at 4 ^°^C for 1.5 hr. The beads were washed three times with 500 µl of lysis buffer without PMSF and three times with 500 µl of phosphatase buffer (50 mM Tris-HCl pH 7.5, 5% glycerol, 0.2% Tween 20, 0.5 mM DTT). The beads were incubated with 20 µl of phosphatase buffer with 1 µl ^32^P-MBP for 10 min at 30ºC gently agitated every few minutes to allow for mixing of the beads.10 µl of sample buffer 4X was added and samples were incubated in a boiling water bath for 5 min and resolved on a 9% SDS-polyacrylamide gel. ^32^P released from phosphorylated MBP migrates at the dye front and was quantified using a Typhoon 9410 Phospho Imager and Image J. The amount of phosphatase substrate was not substantially depleted during the reaction, so substrate was present in excess.

### Microscopy

Photographs of yeast cells were taken using an Zeiss-Axioskop2 Plus microscope fitted with a x63 Plan-Apochromat 1.4NA objective and an AxioCam HRm camera (Carl Zeiss). Images were acquired using AxioVision software and processed using Image J (Schneider *et al.* 2012).

## Acknowledgments

We thank members of the laboratory for advice and support. We also thank Ted Powers (UC Davis) for the Ypk-pT662 phosphospecific antibody and John Tamkun (UC Santa Cruz) for 12CA5 antibody. This work was supported by NIH grant GM053959.

---

**Supplementary Figure 1:**
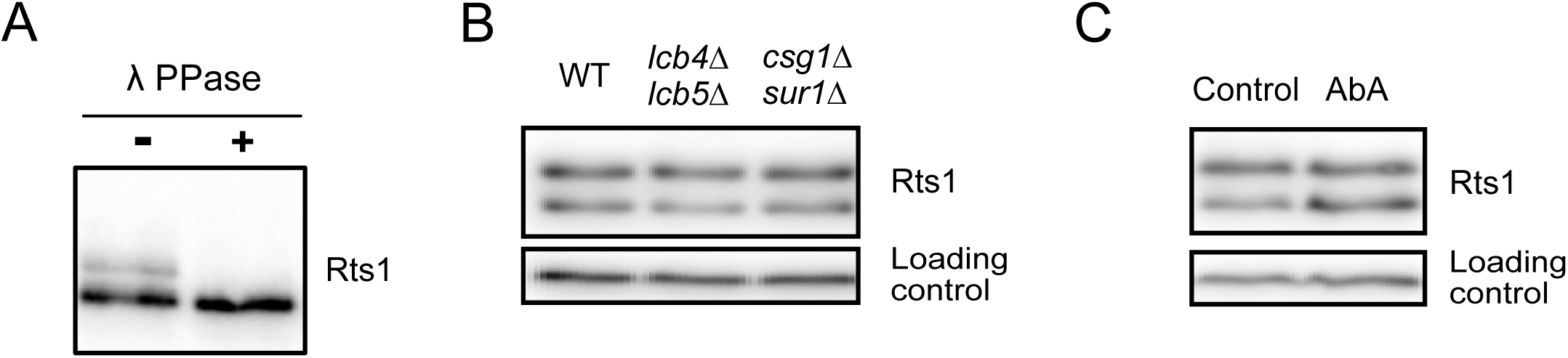
(A) Rts1 is a phosphoprotein. Wild type cells were grown to log phase in YPD medium. A cell extract was made and split into two aliquots. Lambda phosphatase was added to one aliquot. Rts1 was detected by western blot. (B) Rts1 phosphorylation is not affected by other known sphingolipid processing steps. Cells of the indicated genotypes were grown to log phase and Rts1 was detected by western blot. (C) Wildtype cells were grown in YPD for 2h in the presence or absence of 0.5 µg/ml Aureobasidin A. Rts1 was detected by western blot.

**Supplementary Figure 2:**
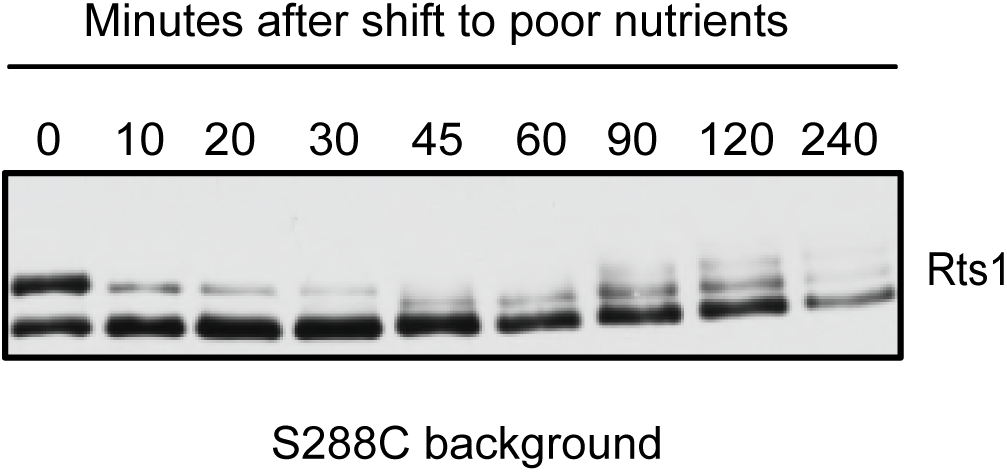
Rts1 responds to a shift from rich to poor carbon differently in the S288C background. Wildtype S288C cells were grown to log phase in YPD medium and were then shifted to YPG/E. Samples were collected at the indicated times and Rts1 phosphorylation was analyzed by western blot. Rts1 in the A364A background showed the same behavior as the W303 background.

**Supplementary Figure 3:**
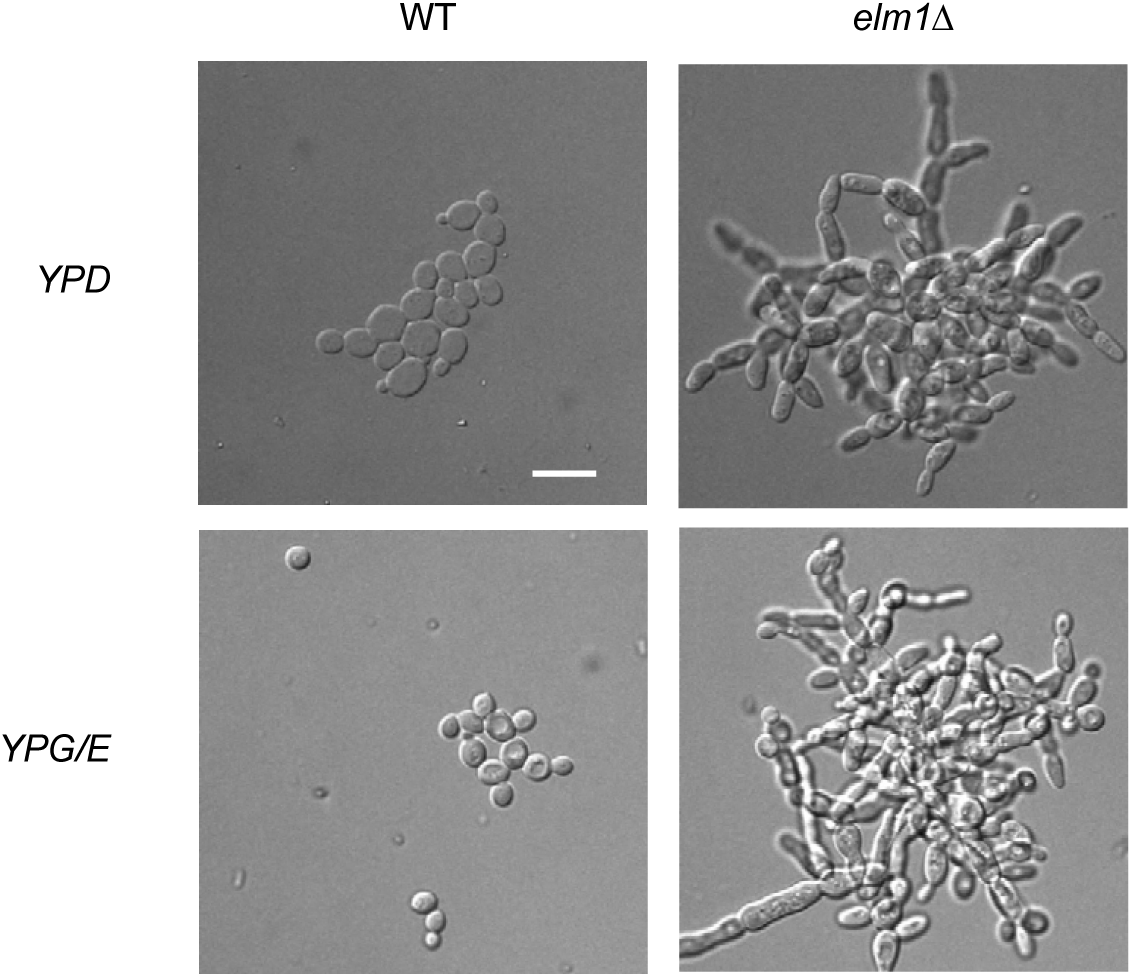
*elm1∆* cells do not reduce their size in poor carbon conditions. Wildtype and *elm1∆* were grown to log phase in YPD or YPG/E medium and photographed. Size scale, 10 µM.

